# Pharmacogenomics Guided Spaceflight: the intersection between space-flown drugs and space genes

**DOI:** 10.1101/2024.01.16.575951

**Authors:** Theodore M. Nelson, Julianna K. Rose, Claire E. Walter, Gresia L. Cervantes-Navarro, Caleb M. Schmidt, Richard Lin, Emma Alexander, Jiang Tao Zheng, Benjamin S. Glicksberg, Julian C. Schmidt, Eliah Overbey, Brinda Rana, Hemal Patel, Michael A. Schmidt, Christopher E. Mason

## Abstract

Ten years ago, it was predicted that the multi-omics revolution would also revolutionize space pharmacogenomics. Current barriers related to the findable, accessible, interoperable, and reproducible use of space-flown pharmaceutical data have contributed to a lack of progress beyond application of earth-based principles. To directly tackle these challenges, we have produced a novel database of all the drugs flown into space, compiled from publicly available ontological and spaceflight-related datasets, to exemplify analyses for describing significant spaceflight-related targets. By focusing on mechanisms perturbed by spaceflight, we have provided a novel avenue for identifying the most relevant changes within the drug absorption, distribution, metabolism, and excretion pathways. We suggest a set of space genes, by necessity limited to available tissue types, that can be expanded and modified based on future tissue-specific and mechanistic-specific high-throughput assays. In sum, we provide the justification and a definitive starting point for pharmacogenomics guided spaceflight as a foundation of precision medicine, which will enable long-term human habitation of the Moon, Mars, and beyond.

**Graphical Abstract:** 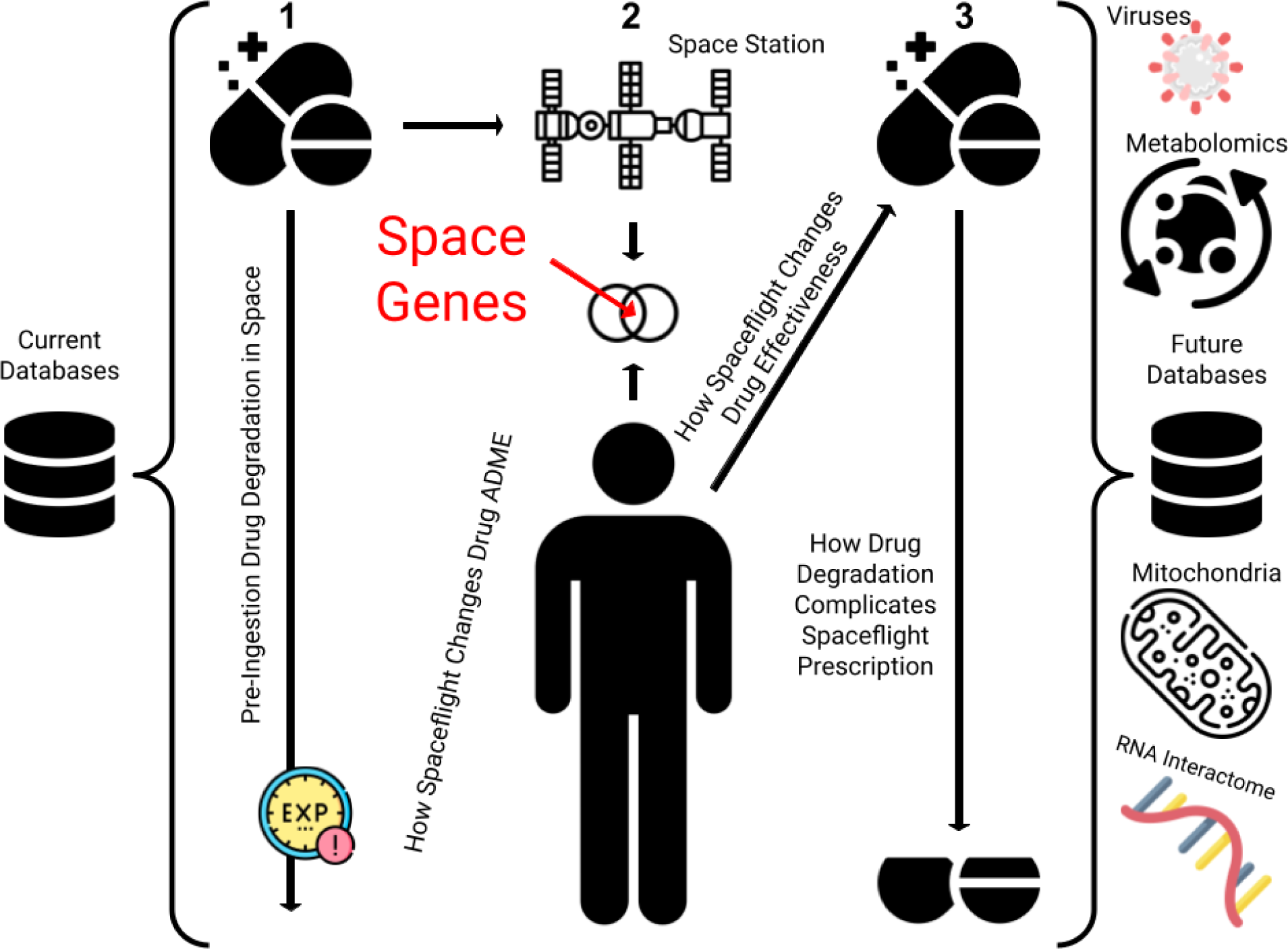

## Introduction

Pharmacogenomics generally refers to the effect of genetic variability on drug efficacy and safety, mediated by drug concentration and mechanism of action.^1,2^ Pharmacogenomics is thereby a fundamental component of precision medicine, defined as the tailoring of medical treatments to the individual characteristics of each patient. Crucially, pharmacogenomics can help predict and protect patients from genetically mediated adverse drug reactions and ineffective drug formulations. For example, individuals with *HLA* allele B*1502, more prominent in the Han Chinese population, are at increased risk for developing Stevens–Johnson syndrome when treated with carbamazepine, a common medication used for neurological conditions like epilepsy.^3^ In terms of drug formulation advisement, individuals with 2 inactive copies of *CYP2D6* exhibit limited metabolism of codeine; these patients are advised to avoid codeine treatments.^4^ Beyond simply advising whether a medication should be prescribed, pharmacogenomics can provide precise dosage recommendations based on individual patient genotypes. For instance, patients taking warfarin, a commonly used blood thinner, require varying dosages of the medication based on *VKORC1* and *CYP* genotype.^5^

Spaceflight-induced suppression of the immune system has led to an examination of potential pharmaceutical remedies, including the application of novel drugs, increasing the risk of polypharmacy within spaceflight environments.^6^ To date, these have not been related to individual genotype profiles.

The field of space pharmacogenomics to date has focused on how to apply earth-based pharmacogenomic principles to clinical practice involving astronauts, both from the perspective of short-term low Earth orbit travelers and future long-term human spaceflight missions.^7^ The implication of current astronaut pharmaceutical usage in space, as well as a clinical perspective regarding the implications of drug metabolizer genotypes in relation to these, are well summarized by Blue et al.^8^ and Schmidt et al.^9^, respectively. While these reviews mention the significant physiological effects of spaceflight, there was insufficient high-throughput data at the time of those writings to tie the individualized spaceflight response to possible downstream pharmaceutical effects.

The previous ten years of human astronaut research has revealed individualized yet wide-ranging transcriptomic effects resulting from both short- and long-term low Earth orbit spaceflight.^10,11^ The development and expansion of the NASA GeneLab multi-omics database,^12^ in combination with the Ames Life Science Data Archive,^13^ have represented a pioneering step in achieving Findable, Accessible, Interoperable, and Reproducible (FAIR) database curation.^14^ Ten years ago, corresponding with the advent of the sequencing revolution, spaceflight-guided personalized medicine was first proposed.^15^ We believe that now is the appropriate time to again address this possibility, especially in the context of spaceflight-guided pharmacogenomics.

In our view, space genes represent a primary driver of these individualized spaceflight transcriptomic responses. These genes are not limited to drug metabolizing genes, but can include all known drug-gene interactions dysregulated by spaceflight. Genotype, epigenetic markers, transcriptomic upregulation or downregulation, post-transcriptional modifications, post-translational tagging, and higher-order interactome complexes are likely complicated by additional spaceflight-related variables, most prominently isolation, microgravity, and space radiation.^15^ Although current spaceflight literature, due to the limited size of astronaut cohorts, generalizes potential intra-person variability to inter-person spaceflight effects, we believe the application of earth-generated knowledge bases can enhance personalized pharmacogenomic treatment in space, especially with the advent of in-flight sequencing technologies.^16,17^ Moreover, the integration of FAIR data processes into future human spaceflight research practice will allow for seamless comparison to better inform clinical recommendations.

Within this framework, we will describe current field-specific databases related to spaceflight pharmaceuticals, introduce our own expanded knowledgebase, find space genes, and hypothesize the implications of these annotations on three critical drug processes: stability, metabolism, and efficacy/safety. Lastly, we will speculate on future possibilities within pharmacogenomics multi-omics integrative analysis.

### Spaceflight-related Data Curation Practices

Space-related research possesses a high barrier to entry, due to limited spaceflight opportunities, data availability, and privacy concerns, resulting in an increased frequency of interdisciplinary collaborations, and decreased statistical power within studied patient cohorts.^18^ Due to these limitations, a digital ecosystem of databases has emerged to preserve information in a FAIR manner.^14^ Database curation and management are maintained by relevant space agencies and supported by international cohorts of subject-matter experts. The aim of these platforms is to discover spaceflight-induced phenotypes across astronaut missions and research studies, reducing future biological threats to long-term human spaceflight through assessment and targeted application of countermeasures.

Even though these databases have been established to produce valuable insights from previous missions, the input is sparse. A few notable studies within the previous decade include a comprehensive examination of the astronaut microbiome,^19^ along with the NASA twins study, which provided a paired-ground spaceflight experiment, although limited to one replicate, analyzing changes pre-, in- and post-flight.^10^ Even though these provided more comprehensive portraits of baseline physiological effects, they did not extensively characterize spaceflight pharmacological-transcriptomic interactions.

We next describe three field-specific databases in terms of their capacity for FAIR data integration: the *NASA Technical Report Server* (https://ntrs.nasa.gov/), the *NASA GeneLab* database (https://genelab.nasa.gov/), and the *Space Omics and Medical Atlas* (https://soma.weill.cornell.edu/) (SOMA).

The first of these, the *NASA Technical Report Server*, is a collection of NASA-affiliated or funded datasets, conference publications, and scientific articles. Although expansive in breadth, the diversity of source material makes the information difficult to integrate.

In contrast, the *NASA GeneLab* repository focuses on a narrower set of data types related to multi-omics sequencing, and allows for uniform processing and integration. This platform is now integrated with the Ames Life Sciences Data Archive (ALSDA). First published in 1994, these combined platforms contain a diverse array of population, microscopy, transcriptomic, genetic, metabolomic, and proteomic data. Experiments have been conducted in a wide variety of organisms as well as microgravity simulation systems. Notably, more than 375 rodent studies have been carried out to understand the mechanisms that affect organisms in conditions ranging from radiation exposure to microgravity treatments. In terms of actual spaceflight studies, these repositories contain over 700 experiments conducted aboard the International Space Station (ISS), the NASA/MIR space station, Bion/Cosmos, Gemini, Biosatellites, Apollo, Skylab, and the Russian Foton missions.

The *Space Omics for Microgravity Atlas* (SOMA), hosted by Weill Cornell Medicine’s Epigenomics Core Facility, is a specialized platform designed to facilitate the analysis of omics data related to space research. It currently houses data from the Inspiration4 mission. Ultimately, study-specific databases such as the *SOMA* Browser, which catalogs the data from SpaceX’s civilian missions, can provide an even deeper layer of integration and visualization support, among diverse and occasionally novel sample collection profiles.

When considering these three databases, the first is difficult to summarize, the second implements FAIR data principles, and the third provides specialized support and immediate results. The collection of current spaceflight pharmacogenomics knowledge mirrors the first of these databases. Ultimately, technical descriptions of drug use are collected within many disparate literature reviews, from studies which collected different kinds of measurements in order to monitor astronaut health. Although generalist sequencing repositories such as the European Nucleotide Archive (https://www.ebi.ac.uk/ena/browser/) have implemented metadata tags in order to better track these kinds of conditions, current NASA-specific repositories do not provide extensive health information or metadata tags to protect the privacy of their astronauts.

### Cataloging Government-Approved Drugs in Space

Given the lack of FAIR data curation within this field, we began this project with the objective of building a comprehensive drug catalog. As there was limited evidence regarding spaceflight-induced alterations to government-approved drugs and their metabolism, agencies built space medicine kits to support the anticipated needs of astronauts during missions. The ISS is stocked with two types of “Med Kits”, *Convenience* and *Contingency Kits*, together containing about 200 pharmaceuticals commonly used on earth. These drugs were flown along with the astronauts on many missions, summarized in publications related to either the Space Shuttle missions or International Space Station missions.^20,21^ The most up-to-date medication usage data was obtained from the “Dose Tracker” study. The Dose Tracker study goals were to collect in-mission medication use data and to inform questions regarding spaceflight-associated changes in pharmacokinetics or pharmacodynamics. The study utilized an iOS app to collect medication name, dose, dosing frequency, indication, perceived efficacy, and side effects on six volunteer ISS crew members and five crew members on the ground. The study reported an average of 20.6 medication use entries per subject per flight week.^22^ Beyond the Dose Tracker Study and individual reviews, few comprehensive databases exist for cataloging drug use in space.^8^

In order to more comprehensively address this challenge, nonprofit BioAstra, Inc (https://www.bioastra.org/) hosted a space medicine mining competition with the aim of cataloging drug usage in spaceflight environments. Ultimately, fifty unique publications were parsed, supplying 394 table entries, with each unique drug mentioned within a single publication serving as the minimum basis for a row (**Table 1**). These studies were sourced from diverse disciplines, examining drug effects on both prokaryotic and eukaryotic organisms.

**Table 1:** Catalog of Drug Usage within Spaceflight Environments.

The compiled database contains 218 unique drugs and drug combinations. These medications were sometimes tested in tandem; examples include scopolamine-dextroamphetamine and zolpidem-zaleplon. Of the medications listed within this database, the medications most mentioned within research were zolpidem (12), ampicillin (10), erythromycin (10), gentamicin (10), acetaminophen (8), promethazine (8), and tetracycline (8).

Of the 218 pharmaceuticals listed on the database, 45 of them are currently employed as part of the ISS medical kit, as listed in the 2016 NASA Emergency Medical Procedures Manual for the International Space Station (https://www.governmentattic.org/19docs/NASA-ISSmedicalEmergManual_2016.pdf). These medications include some of the more common medications as per the compiled database, including zolpidem, erythromycin, acetaminophen, promethazine, and tetracycline. These 45 medications are in relation to the 85 listed medications available on the ISS, demonstrating that there are a significant number of medications currently aboard that have not been addressed within current spaceflight-pharmacology literature.

Notably, when comparing the ISS medical kit components and our review of published literature, four combinations of drugs listed as part of the medical kit were not explicitly referenced within our database. These pairs include: sulfamethoxazole and trimethoprim, hydrocodone and acetaminophen, ciprofloxacin and dexamethasone, and tobramycin and dexamethasone (https://www.governmentattic.org/19docs/NASA-ISSmedicalEmergManual_2016.pdf). While each of these medications was listed on the compiled database individually, there were no studies found which addressed the potential effects of coupling these medications in-flight. The application of FAIR data reporting standards in future spaceflight missions is crucial to preventing these discrepancies within clinical records.

### Abiotic Stability of Government-Approved Drugs in Space

Due to these discrepancies within reporting standards, as well as the importance of accurate prescriptive treatment of astronauts, a brief discussion on drug degradation and expiration during spaceflight in an abiotic context is warranted. To assess drug storage specifically, studies have most often examined the percentage of active pharmaceutical ingredients (API). Particularly striking are results indicating that medications flown aboard the ISS, including levothyroxine, promethazine, dextroamphetamine, and ciprofloxacin, did not meet regulatory requirements in terms of API content prior to their expiration date.^23^ These medications, particularly the latter three, are highly light sensitive, and therefore likely saw significant increases in their degradation in space.^23^ In a separate study, API degradation and impurity were observed for some of the most commonly utilized drugs, including aspirin, ibuprofen, loratadine, modafinil, and zolpidem after storage aboard the ISS for 550 days.^21^ Similar results have been shown for scopolamine, one of the most heavily prescribed drugs for astronauts,^21,24–27^ specifically to treat space adaptation syndrome and space motion sickness.^28^ Similar studies have been further summarized in Blue et. al., supporting a similar conclusion.^8^

Nevertheless, it should be acknowledged that the observed changes in API are relatively small, suggesting that this ultimately may not contribute to clinical outcomes.^29^ Notably, all of the changes in drug stability aboard the ISS were within 10% of the changes seen at the Johnson Space Center.^29^ These differences may be the result of confounding variables, including drug formulation and pre-flight storage conditions. In a study of different Vitamin B formulations within multivitamin brands, it was shown that the API of Vitamin B_1_ was changed significantly in one brand after storage on the ISS for 12 or 19 months, while the second brand demonstrated no statistically significant changes.^30^ Further documentation and studies are necessary to both resolve this question, and evaluate downstream clinical effects.

### Defining the Biological Foundations Associated with Government-Approved Drugs Used in Space

As discussed in the Introduction, genes constitute the foundational unit of biological analysis, currently employed in transcriptomic and proteomic analyses. These transcriptomic and proteomic states therefore represent a complete cellular portrait for a particular physiological context. Within this review, we take a novel approach to defining a set of related space genes, utilizing our drug database as a starting point. To define a set of related genes impacted by spaceflight-flown pharmaceuticals, we utilized the list of drugs known to have been applied in space, along with the drug-gene interaction database DGIdb (https://www.dgidb.org/) in order to generate a gene set potentially affected by in-flight drug usage. After collating the results from both database versions, we found that out of the 218 unique drugs within our space medicine database, 190 had interactions with 772 unique genes, for a total of 2,318 interactions (**Table 2**).

**Table 2:** Spaceflight-Flown Drug-Gene Interaction Database.

The newer DGIdb database version ranks drug-gene interactions via a scoring metric. This is collated based on evidence scores, related to the number of supporting publications and sources, the ratio of average known gene partners for all drugs to the known partners for the given drug, and the ratio of average known drug partners for all genes to the known partners for the given gene, according to https://www.dgidb.org/. We utilize a lenient cutoff of `.2` for discussion purposes within this review. 681 of the ranked interactions pass this cutoff, while 377 interactions do not have an interaction score assigned (**Figure 1**).

**Figure 1.**
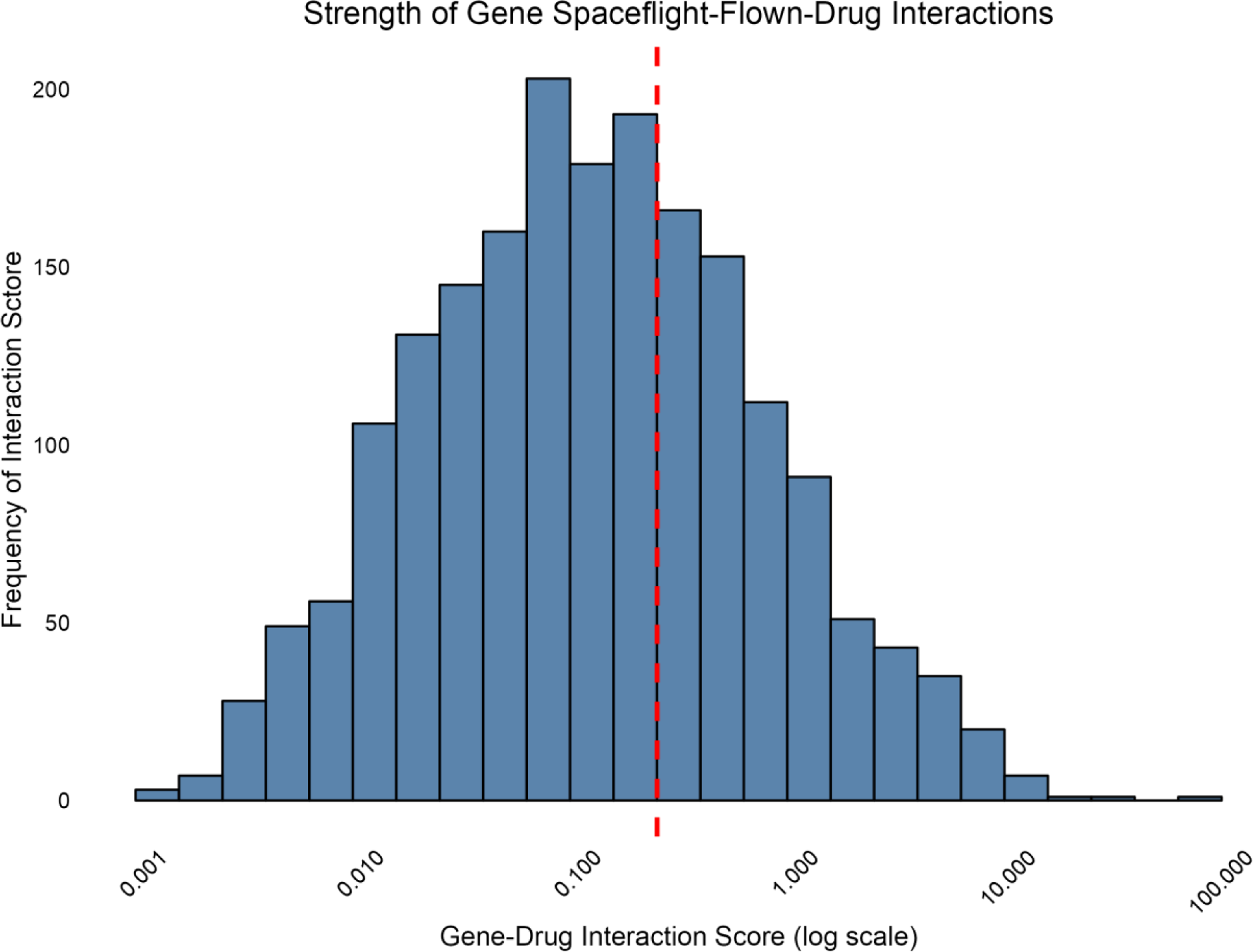
Strength of Gene Spaceflight-Flown-Drug Interactions with a line at the border demarcating an interaction strength of .2 [note the log scale on the x-axis].

Unsurprisingly, these genes are as a collection most associated with their respective responses to different chemicals (**Supplemental Figure 1**), further indicating their important role within the mechanistic action of the selected drugs. Downstream effects could result from many different potential mechanisms related to these genes.

### Hypothesizing Spaceflight-perturbed Drugs with High-throughput Proteomics

The genes mentioned above represent important targets for these common drugs, although their specific relevance to the spaceflight environment has not been established. To investigate this relevance, researchers often examine changes within gene expression across conditions, which represents a classical mode of defining gene ontologies.^31^

When cells from the Caco-2 cell line, a human male colorectal adenocarcinoma, were treated with simulated microgravity conditions, researchers found a host of proteins with differential abundances at both 48 and 72 hours of microgravity using label-free shotgun proteomics.^32^ Capturing a total of 6,109 proteins with this assay, the authors found 62 with differential abundance.^32^ These include decreased basal activation of *NFκB*, a transcription factor relating to inflammatory responses, underexpression of *MTFR2* and *MT-ATP8*, proteins related to ATP synthesis and mitochondrial functions, *CDH17*, a calcium dependent cell adhesion protein, *CDK2*, and many other proteins related to intestine morphology and peptide transport.^32^ Certain overexpressed proteins, including *HNRNPDL*, which has been implicated as an oncogene, upregulated in pancreatic tumor cells under the study’s simulated microgravity conditions, suggesting potential confounding results from the cancer cell model system.^32^

When comparing this proteomic set to the 772 genes associated with space-flown medicines, there was a total overlap of three genes: *APOA1*, interacting with furosemide and progestin, *CDH17*, interacting with dexamethasone, and *DPEP1*, interacting with cilastatin and dexamethasone.

### Hypothesizing Spaceflight-perturbed Drugs with High-throughput Transcriptomics

A convergent approach defines spaceflight-perturbed genes utilizing RNA-sequencing assays, which capture gene expression across conditions.^33^ For this purpose, we selected the closest public analogue of human spaceflight available within the NASA GeneLab repository; namely, a study of human induced pluripotent stem cell-derived cardiomyocytes, which utilized a well-controlled ISS spaceflight perturbation, as opposed to microgravity simulation devices.^34^ Although cell types specific to drug metabolizing cells within the liver would have been preferable, we selected the most relevant dataset available for this application.

To indicate drugs whose mechanistic actions may be perturbed by spaceflight, we intersected the set of 772 unique genes associated with spaceflight-applied drugs with the set of differentially expressed genes, available within the supplemental materials of Wnorowski et. al.^34^ There were 48 genes whose mechanistic perturbation intersected with current spaceflight-flown drug associated genes: *ABCA1*, *ABL1*, *ADRB3*, *APOE*, *ATF4*, *B4GALT2*, *BAX*, *CHRM3*, *CREBBP*, *CYP1B1*, *CYP2D6*, *CYP3A43*, *ERCC1*, *FOS*, *FTO*, *GCLC*, *GGT1*, *GLP1R*, *GNAS*, *GSR*, *GSTP1*, *GTF2B*, *HRH1*, *HSPA5*, *HSPA8*, *IMPDH1*, *JUN*, *JUNB*, *MECP2*, *MVK*, *PIK3R2*, *PLOD1*, PRDX1, *PTH1R*, *RAD52*, *RPS19*, *SCN9A*, *SDHB*, *SLC19A1*, *SLC23A2*, *SLC29A2*, *SOD1*, *STS*, *TGM2*, *TMEM167A*, *UGT1A4*, *WRN*, and *ZBTB22*. The gene ontology for each of these genes is described in Supplemental Table 1.

Collectively these influenced the following 36 drugs: acyclovir, alendronate sodium, ascorbic acid, aspirin, atorvastatin calcium trihydrate, brompheniramine maleate, cefotaxime sodium, cetirizine hydrochloride, chlorzoxazone, cinnarizine, ciprofloxacin, copper chloride, cyclizine, dexamethasone, dimenhydrinate, diphenhydramine, doxycycline anhydrous, fluorouracil, gentamicin, hydrogen peroxide, levofloxacin anhydrous, melatonin, mercaptopurine, metoclopramide hydrochloride, ofloxacin, ondansetron hydrochloride, progestin, quetiapine fumarate, selenomethionine, sulfamethoxazole, tetracycline, thrombin, triprolidine, vitamin d, vitamin e.

When comparing between the differential sets associated with either the gene-drug interaction database, the proteomic gene set, and the transcriptomic gene set, we find no overlap between any of the indicated space genes (**Figure 2**). As such, we recognize the preliminary nature of these predictions. In the remainder of this review, we offer two avenues of interpretation; the first relying on the strength of the interaction within the predicted targets as a measure of importance and the second contextualizing these results with a focus on previous literature within spaceflight pharmacogenomics. For example, we provide commentary on the *CYP* family, which has significant known functions within drug metabolism,^9^ while also speculating on genes such as *MVK*, which has not been previously studied in the context of spaceflight.

**Figure 2.**
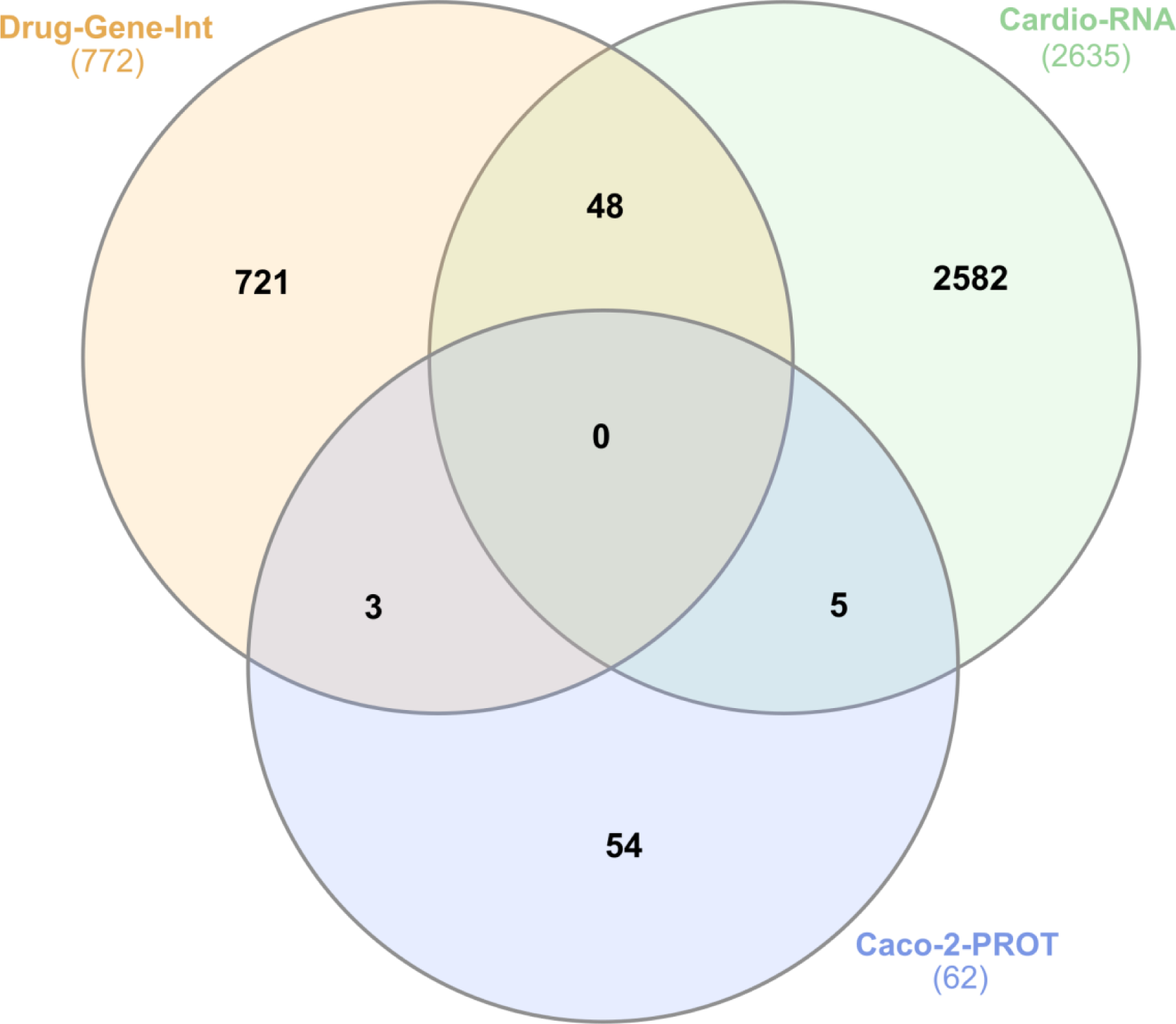
Intersection between published spaceflight-flown drug-gene interactions, differentially expressed gene sets from ISS-flown cardiomyocytes, and differentially expressed protein sets from simulated-microgravity Caco-2 cells. Chart produced with Interactivenn.^99^

### Biotic Stability of Government-Approved Drugs in Space

Whereas we previously examined drug storage stability within an abiotic context, once ingested during spaceflight the drug encounters a myriad of new biologically-related challenges to carry out its function. We break these into three separate stages: biotic stability, describing drug processing pre-target; effectiveness, characterizing the interaction with specific target genes and proteins; and degradation, relating downstream chemical degradation products leftover from drug usage. To further distinguish between the elements of biotic stability, we examine individually each stage of absorption, distribution, metabolism, and excretion (ADME) and how they differ within a spaceflight environment through selected examples (**Figure 3**).

**Figure 3.**
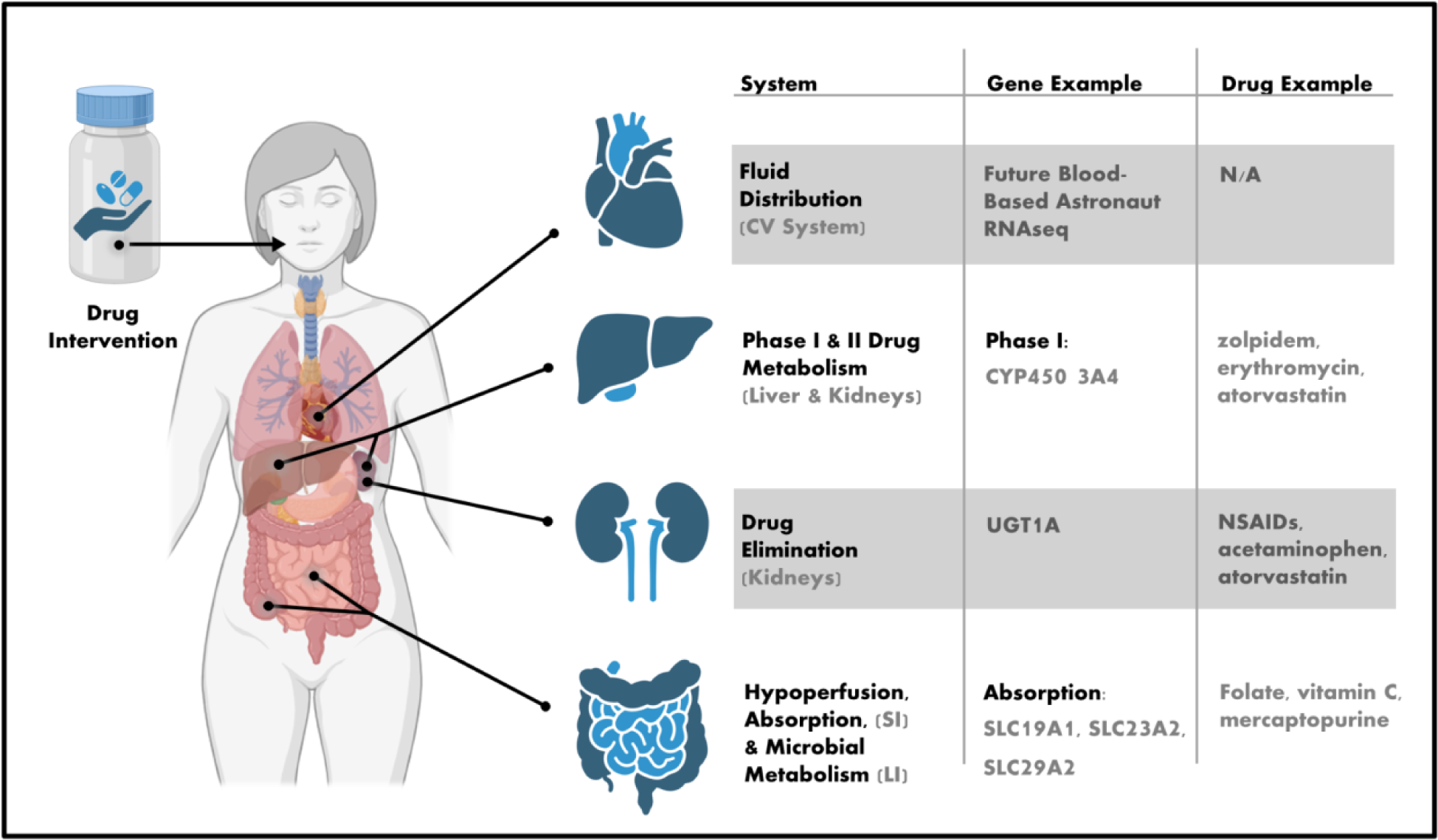
ADME model of Altered Drug Metabolism in Space. (CV: cardiovascular, SI: small intestine, LI: large intestine)

Orally ingested drugs, as opposed to injected drugs, are initially subject to: 1) increased drug breakdown, due to delayed gastric emptying and changes in microbial community structure and 2) reduced drug absorption, due to accelerated intestinal transit, fluid shift-associated hypoperfusion of the GI tract, and changes in expression of GI enzymes and transporters.^28^ As an example, we observed three solute transporter genes, *SLC19A1*, *SLC23A2*, *SLC29A2*, for folate, Vitamin C, and nucleosides, respectively within our set of space genes; in the case of *SLC23A2*, functional disruption has been linked to gastric cancer.^35^ Both *SLC19A1*, *SLC23A2* demonstrated selective spaceflight-related downregulation, whereas *SLC29A2* was upregulated.

Once absorbed, the spaceflight environment significantly alters drug distribution for orally ingested as well as injected drugs. These factors are myriad and individual response is variable, relating to the following: 1) fluid shift: the headward fluid shift increases natriuresis and diuresis, impaired lymph flow and lymphatic drainage,^36^ decreases thirst, and increases evaporation through the skin and lungs, 2) fluid redistribution from plasma to extracellular volume; and thereafter from extracellular volume to intracellular volume, 3) decreased volume, decreased drug distribution, and increased plasma concentration, 4) altered binding expression and plasma concentration, 5) endothelial dysfunction, and 6) changes in hepatic blood flow.^28^ Although our produced gene set does not necessarily capture hepatic shifts well, there are opportunities within this field, especially based on the information available via the SOMA Browser (https://soma.weill.cornell.edu/), to compare cell-free RNA profiles and blood cell transcriptomic profiles from recent missions of NASA and SpaceX astronauts.^10,11^

Having examined absorption and distribution changes, we turn to drug metabolism, which is affected by: 1) expression and, thus, enzyme concentration changes of the phase I and phase II drug metabolism system, 2) pharmacometabolomic signatures associated with a certain drug intervention or metabolic state as it relates to drug metabolism, and 3) food components and other xenobiotics as a result of polypharmacy and other environmental exposures that are also known to be inducers and inhibitors of the Cytochrome P450 (CYP450) pathway.^9^ Within our set of space genes, we observed differential expression of *CYP1B1, CYP2D6, CYP3A43*, which are all major components of the CYP450 pathway, a superfamily of monooxygenase enzymes that perform various oxidation reactions; these enzymes and their respective reactions are the dominant metabolic process in phase 1 metabolism, occur mostly within the hepatocytes of the liver and are characterized by oxidation, reduction, and hydrolysis reactions.^37^ Pharmaceutical inhibition or induction of CYP450 enzymes is the leading cause of drug-drug interactions, as the pharmaceutical inhibition or induction of one enzyme can inhibit the metabolism of another drug by the same enzyme.^38^ By this mechanism, in the case that the volume or function of primary CYP450 enzymes was altered significantly, it could have detrimental effects on the metabolism of certain drugs. CYP450 enzymes also play a major role in the conversion of prodrugs to active metabolites and altered function due to gene expression changes or functional variants leads to adverse effect or poor drug response in as in the case of *CYP2D6* in the conversion of codeine to morphine^39^ and *CYP2C19* mediated conversion of clopidogrel.^40^ Within cardiomyocytes, *CYP1B1* and *CYP2D6* demonstrated downregulation during spaceflight, whereas *CYP3A43* was upregulated.

Multiple studies have similarly observed changes in CYP450 expression during spaceflight.^41–43^ In a convergence with proteomic activity assays, one specific study showed a decrease in CYP450 activity by 50%.^28,44^ Although these studies did not address the possibility of decreased pharmaceutical metabolization by CYP450, we believe that this is an important key consideration for pharmacogenomics. Future studies should determine whether the CYP450 enzymes are similarly affected in human astronauts - and the downstream implications for drug metabolism.

Liver homeostasis relies in part on glutathione, an important compound for protecting against oxidative stress.^45^ Glutathione additionally serves as a crucial molecule in drug conjugation.^46^ We noted *GCLC* and *GGT1* upregulation in the cardiomyocyte transcriptomic spaceflight profiles, important for glutathione production, suggesting an increased response to oxidative stress.^47,48^ These results suggest convergence with current hypothesis regarding increased levels of oxidative stress within spaceflight environments.^49^

We postpone our discussion of how physiological and gene expression changes alter elimination through hepatic metabolism and decreased urinary output to the Degradation and Excretion of Drug Products section.

### Effectiveness of Government-Approved Drugs in Space

Separate from questions of drug absorption and processing is whether the mechanistic action with the primary target of interest is disrupted during spaceflight. In particular, the example of promethazine was particularly extreme from a clinical perspective. Pharmacodynamics were significantly altered on space shuttle flights, with 4.8% of astronauts experiencing sedation upon administration of this drug in comparison to the 60-73% who experienced sedation in ground controls.^50^ Based on the drug-gene interaction database, it might be suggested that this is due to an interaction with either *CHRM4* or *CYP1A1*. Nevertheless, we found no evidence within the cardiomyocyte transcriptomic or Caco-2 proteomic profiles that these genes were affected by spaceflight. Given these drastic effects, predicting potential loss-of-function effects may be accomplished from a mechanistic perspective, better yet utilizing hepatic cell-type specific tissue. By examining changes within the genome, transcriptome, epitranscriptome, and proteome, potentially ineffective mechanisms of action can be avoided.

Instead of relying on clinical spaceflight observations, we can also select the strongest interaction partners and predict potential mechanistic interruptions. To demonstrate this, we have selected interaction with the greatest score (13.10667864), between the *MVK* gene and sodium alendronate (**Table 2**). The human *MVK* gene encodes for mevalonate kinase enzyme (MK), the second enzyme in the mevalonate, or isoprenoid, pathway, oftentimes considered the “bottleneck” of the pathway.^51–53^ The mevalonate pathway (MVP) functions in synthesizing sterol and non-sterol isoprenoids including heme A, ubiquinone, and cholesterol, deeming it vital to numerous bodily processes.^51^ The larger MK pathway is targeted by statins and bisphosphonates for cholesterol management, cardiovascular health, and more recently cancer treatment.^54^

However, MK itself is currently linked only with sodium alendronate, a bisphosphonate derivative, in the context of treating mevalonate kinase deficiency (MVD). MK is controlled by a negative feedback loop via geranyl and farnesylpyrophosphate, two intermediates towards the end of the MVP, insinuating that MK could be indirectly targeted by changing concentrations of these intermediates.^53^ In the case of a 14 year old boy with MVD, sodium alendronate was used to completely reverse the symptoms of the disease. The mechanism, however, is unknown, and the clinicians in the study expressed surprise that the treatment was effective at all.^55^ Sodium alendronate is more commonly used to relieve symptoms of osteoporosis despite significant recorded side effects.^56^ Therefore, sodium alendronate has been a drug of interest for the prevention of bone loss in astronauts.^57,58^

While more studies are needed to elucidate the underlying link between MK and the effect of sodium alendronate, within cardiomyocytes, *MVK* is upregulated in spaceflight. As a result, we can hypothesize that this interaction could have different pharmacokinetics within the spaceflight environment. Integrating further pharmacogenomic resources will allow better clinical recommendations regarding this drug, as well as determining whether the change in gene expression is based on specific regulation of the *MVK*, or upstream effects related to changes within the broader pathway. These two approaches to evaluating drug effectiveness, from a first-principles perspective of either gene expression or clinical presentation, have the potential to converge within a FAIR curated knowledge base repository.

### Degradation and Excretion of Drug Products

Returning to the final piece of the model described in **Figure 3**, once drugs have affected their primary function, they are degraded and excreted. Similar gene-based mechanisms as modeled above can, from a biological perspective, alter elimination through hepatic metabolism and decreased urinary output. Within the cardiomyocyte transcriptome sequencing data, we note that a key component of the glucuronidation pathway, *UGT1A4*, was upregulated in flight, allowing potentially for more efficient waste elimination.^59,60^ Interestingly, the FDA recommends testing for a genetic variant (in/del) causing reduced expression another *UGT* gene, *UGT1A1,* before the administration of irinotecan due to elevated risk for neutropenia in patients homozygous for the reduced expression variant *28.^61^ Nevertheless, with regard to spaceflight-specific effects of degradation of drugs within *in vivo* biological systems, there is a severe lack of data within public literature.

Therefore, in this context, we instead examine the possible effects of abiotic degradation, as described in the section **Abiotic Stability of Government-Approved Drugs in Space**, on the biological systems they encounter. Nevertheless, these are still difficult to characterize. In a Raman spectroscopy study of drug degradation products in an abiotic context pre-ingestion on the ISS, it was found that some degradation products, specifically azaerythromycin A and norepinephrine, do not exhibit significant spectral differences from their API counterparts, azithromycin and epinephrine, respectively.^62^ Notably, novel degradation products have not been identified within abiotic processes within studies of commonly used space pharmaceuticals, including aspirin, ibuprofen, loratadine, and zolpidem.^21^ Nevertheless, we briefly review in the following sections the main abiotic degradation products for the most commonly utilized drugs in space.

Zolpidem, the drug that was most commonly used aboard the ISS based upon our database, has four primary degradation products that have been studied: zolpacid, oxozolpidem, zolpaldehyde, and zolpyridine. These compounds were measured after photolysis and oxidation, two methods based on liquid formulations of zolpidem.^63^ While many of these zolpidem degradation products have the same basic structure as zolpidem, which consists of an imidazopyridine ring,^63^ they may have different toxicity, bioavailability, and therapeutic effects than zolpidem.^64^ For example, while few studies have investigated the effects of zolpacid on human health, it has been seen to have a shorter retention time, approximately 39% of zolpidem, indicating that upon degradation, this drug will be less effective at treating astronauts.^65^ Not only that, but the degradation products of zolpidem have highly variable polarity; zolpacid is highly polar due to carboxylic groups, and zolpaldehyde is highly nonpolar.^64^ It is important to note, however, that zolpidem has been deemed stable in its solid form, with degradation occurring more often in solution.^63^ Thus, the degradation products from this drug may not be as much of a concern as some other compounds aboard the ISS. Yet, there is still some indication of degradation of solid zolpidem under the stressors of heat and visible light to around 15% degradation.^66^

The second most significant medication, as determined via the comparison between our compiled database and the current ISS medical kit, was erythromycin. As erythromycin degrades, it is converted into its metabolite ERY-H_2_O through dehydration of the compound.^67,68^ While ERY-H_2_O has primarily been studied in an ecological context, particularly with regards to water contamination, ERY-H_2_O contributes to bacterial erythromycin resistance despite the degradation of the compound itself.^68,69^ This leads to declining drug efficacy, which would be problematic if this medication were to be brought along longer-duration space missions.

Regarding both previous drugs, there are also potential interactome interactions between these effector complexes. Zolpidem is known to rely on the *CYP3A4* gene for the majority of its metabolism.^70^ Erythromycin itself is an inhibitor of *CYP3A4*, suggesting possible delayed effects if these drugs were given together.^71^ These kinds of interactions within compound polypharmacy should also be considered in future space medicine prescription.

Acetaminophen is a drug with several studied impurities and degradation products that have the potential to negatively impact human health, indicating the importance of understanding spaceflight-related drug metabolism and degradation. Prominent degradation products include p-aminophenol and hydroquinone. Hydroquinone is a key product of study, being a benzenediol compound with debated human health impacts.^72^ More specifically, the carcinogenic nature of hydroquinone has been contested, mainly due to its use as a topical agent for skin lightening.

Most studies of hydroquinone carcinogenicity have been conducted in rat models, such as a 2005 study indicating that hydroquinone can build up in the body, as it is not initially metabolized by the liver, which can increase the abundance of carcinogenic metabolites and lead to nephrotoxicity.^73^ However, a variety of human studies have not found significant correlation between hydroquinone and cancer.^74^ With such debates about the human health risks associated with this compound, it is important to be aware of the degradation of acetaminophen to hydroquinone aboard the space station or future long duration space vessels. Furthermore, the EPA lists a number of noncancerous effects such as increased tinnitus, nausea, vomiting, and abdominal cramps upon ingestion of this compound (https://www.epa.gov/sites/default/files/2016-09/documents/hydroquinone.pdf). As such, monitoring acetaminophen degradation in space will be increasingly important as flight durations increase.

Furthermore, promethazine degradation products have been studied, primarily using oxidative degradation methods. These products include formaldehyde, acetaldehyde, and dimethylamine, 10-methylphenothiazine, phenothiazine, 3H-phenothiazine-3-one, phenothiazine 5-oxide, promethazine 5-oxide, and 7-hydroxy-3H-phenothiazine-3-one.^75^ Such products will be important to consider and monitor along long duration space missions. For example, formaldehyde has been implicated in Fanconi Anemia and presents potential impacts on genome instability upon being produced during cellular metabolism.^76^ Not only that, but ingestion of this compound in its liquid state has been associated with numerous health conditions in humans, including metabolic acidosis, proteinuria, abdominal pain, nausea, and dizziness.^77^ While degradation may not be driven by thermal oxidation or promethazine in solution aboard the ISS, the degradation products that have been shown via this methodology should continue to be examined when considering medication for space missions.

The last drug that will be analyzed in depth in this section is tetracycline, a broad-spectrum antibiotic.^78^ The main degradation products of tetracycline are 5a,6-anhydrotetracycline hydrochloride (ATC), 4-epitetracycline hydrochloride (ETC), and 4-epi-anhydro-tetracycline hydrochloride (EATC).^79^ ETC and EATC have been shown to have negative impacts on human health; EATC, for example, can induce Fanconi syndrome, a kidney disorder which can lead to heart attack, vomiting, glycosuria, and proteinuria, among other symptoms.^80,81^ With such problematic effects, the degradation products of tetracycline become a human health risk for long duration space missions.

Many of the degradation products have been studied in aquatic environments, where antibiotics have accumulated due to their increased prevalence. Thus, the degradation products arising from these environments may differ from those aboard the ISS or spacecraft for longer duration space missions. However, the fact that known degradation products have the potential to negatively impact human health indicates the need to conduct further studies of drug degradation products and pathways in the space environment to protect astronauts and future space travelers. Ultimately, maintaining 90% potency, a target within Mehta & Bhayani 2017, for medications, likely, remains a future challenge for many classes of drugs within space pharmacogenomics, given the likely variable impacts of long-term spaceflight on drug degradation.^82^

## Conclusion

Within this review, we have evaluated previous database according to FAIR data principles, presented a new database of spaceflight-employed drugs, demonstrated two proof-of-concept definitions for ‘spaceflight response’ based on either high-throughput proteomic or transcriptomic profiles, before emphasizing and introducing components of the pharmacogenomics field — specifically drug stability, metabolism and efficacy/safety - from the perspective of a pharmacogenomics guided spaceflight.^9^

At the present time, the name ‘pharmacogenomics’ may be somewhat of a misnomer, given that this integrative analysis has focused on transcriptomics. Yet, these changes induced by spaceflight are ultimately crucial, especially within the kind of short-term missions that provide data within the field. Ultimately, our vision sees spaceflight pharmacogenomics as encompassing the intersection between every known mechanistic element, capable of being measured via high-throughput analysis, including genetic variation via single-nucleotide polymorphisms (SNPs), which have been implicated in variations to drug response.^83^ Of course, these are crucial to considerations within long-term spaceflight habitation, since radiation, present to higher levels within spaceflight environments, is known to be a major driver of genomic instability.^84^ Across animal, plant, and fungi kingdoms, SNPs associated with spaceflight have been reported.^84–86^ In humans particularly, space radiation left SNPs that were known to be risk alleles for neuro-ocular issues, suggesting a potential functional consequence of these SNPs.^84^ In bacteria, SNPs have been associated with the development of antibiotic resistance.^87,88,89,90^

However, to make evidence-based clinical recommendations, all the above components need to be combined with information regarding drug absorption, distribution, metabolism, and excretion (ADME). The ADME gestalt that begins to form, all things considered, when comparing drug intervention is the spaceflight environment versus its ground-based counterpart, is complex and variable. When considering an individual and their response to orally ingested drugs in space, the compounding variance within a single person can lead to wildly different outcomes intrapersonally and interpersonally, especially when mission duration is considered. This begs for the implementation of an evidence-based, precision drug prescribing paradigm in spaceflight that fulfills several objectives: 1) it understands the totality of drugs given in space and their physiology and omics-based historical behavior, including drug-gene and drug-drug responses 2) it pre-screens all individuals entering the spaceflight environment for features of known variance, including SNP and biomarker profiling, and 3) it prescribes medications according to evidence - based principles, utilizing extant databases and prescribing guidelines. The following sections and material presented in this paper are initially steps aimed at fulfilling objective 1 and standardizing a resource that can be used by the space medicine community. Given the re-emphasis within this paper on genes involved in several known PGx recommendations, we suggest that terrestrial PGx practice, related to SNP profiles, covering approximately 300 drugs, be applied within space medicine preparation and practice.

In addition to a SNP-centric vision, we believe that genomic instability translates to downstream modifications within the transcriptome and proteome. Therefore, it would be prudent to examine a wide range of high-throughput mechanistic outputs, including epitranscriptome modifications,^91^ post-translational modifications,^92^ metabolomics,^93^ chromatin-chromatin interactions,^94^ and RNA interactomes.^95^ Viral diseases remain understudied in space,^96^ along with the role of the mitochondria in spaceflight disease processes.^49^ Additionally, our studies to date are limited based on available data for particular tissue- and transcriptome-types, along with their associated biological gender.^97,98^ Even if meta-analyses are conducted as proposed across current human astronaut studies, they would be generalizing peripheral blood mononuclear cells to be reflective of hepatic transcriptome. Within this review, we have generalized cardiomyocyte spaceflight response as a generalized non-tissue-specific spaceflight response. To incorporate all this information and work past these limitations, there exists a FAIR framework for collecting and storing data: NASA GeneLab.

One limitation of this study regards potential double reporting. Due to confidentiality, the extent to which astronaut drug use was reported in more than one study is not currently known. This risk of ‘double reporting’ does not adversely impact the development of the space gene drug database. However, it may impact the frequency distribution (i.e. the ranking of which drugs are most used in space).

One remedy for the latter is to more carefully identify the individual drugs used by individual astronauts and avoid the double reporting phenomenon. However, this is problematic, as it would violate astronaut confidentiality. Perhaps this latter point could be remedied by a prospective study undertaken by the astronaut office where the frequency distribution can be carefully determined without violation of confidentiality.

In closing, drug-gene interactions will become increasingly significant in the context of civilian spaceflight, where astronauts will exhibit more comorbidities than professional astronauts and will likely be on more medications prior to entering the spaceflight environment. Ultimately, the extent of these impacts (regardless of whether one is a government or civilian astronaut) is based both on the importance of the disrupted mechanism and exposure to the drug. Spaceflight has a major effect on the pharmacokinetics of drugs; a meta-analysis reveals that these effects are rather varied depending on the specific drug class and use case. As such, we have described the most prominent space genes, based on publicly available high-throughput proteomics and transcriptomic analyses, which exemplify the current state of the field.

## Author contributions

TMN, JKR, CEW, RL, JTZ, EGO, CEM conceptualized the study. TMN, JKR, CEW, RL, JTZ, EGO curated the space medicine database. TMN, JKR, CEW, GC wrote the original manuscript draft. TMN, JKR, CEW, GC, EA, EGO, BSG, CMS, JCS, MAS, CEM wrote, reviewed, and edited the manuscript. EGO and CEM administered the project. TMN, CEW, EA, CMS, JCS generated the figures. TMN, JKR, CEW, CMS, JCS, HP, BR, EGO, BSG, MAS, CEM performed final review.

## Acknowledgments

We thank NASA (NNX14AH50G, NNX17AB26G, 80NSSC22K0254, NNH18ZTT001N-FG2, NNX16AO69A, 80NSSC23K0832), and the WorldQuant Foundation.

## Data availability

All relevant data is published in the main tables of this manuscript.

## Code availability

N/A

## Supplemental Figures

**Supplemental Figure 1:**
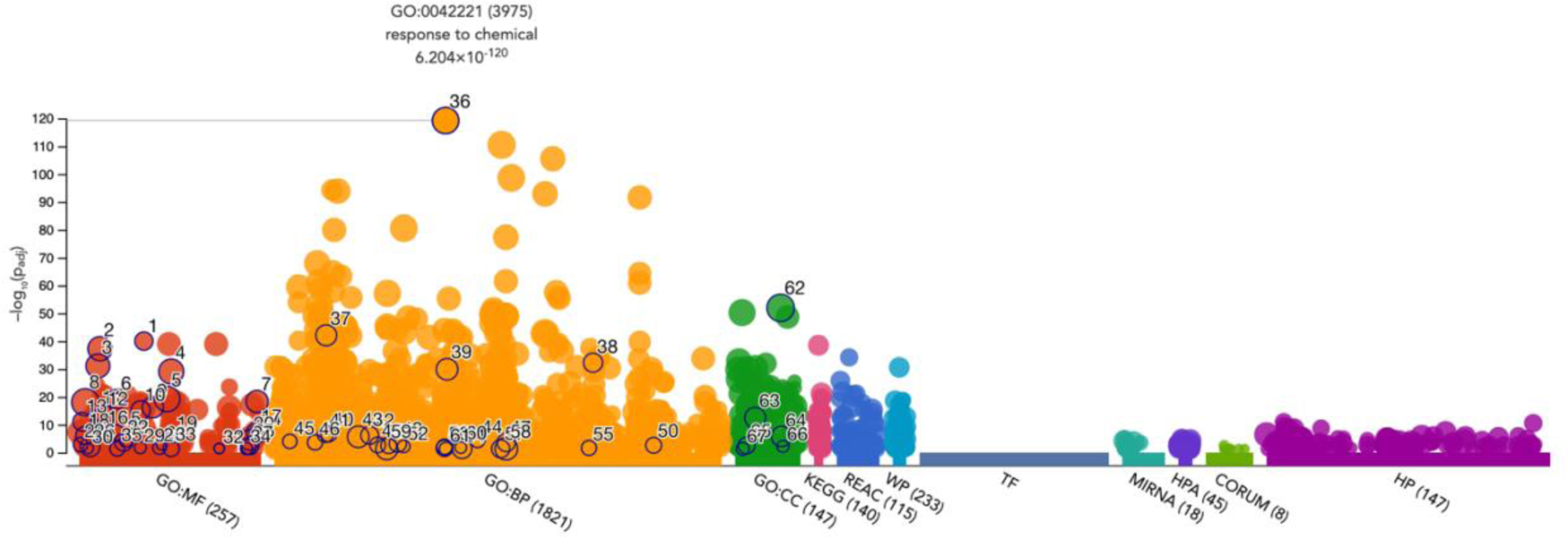
Gene ontology of the 772 unique spaceflight-associated genes demonstrating the largest shift within the term ‘response to chemical.’

## Supplemental Tables

**Supplemental Table 1.**
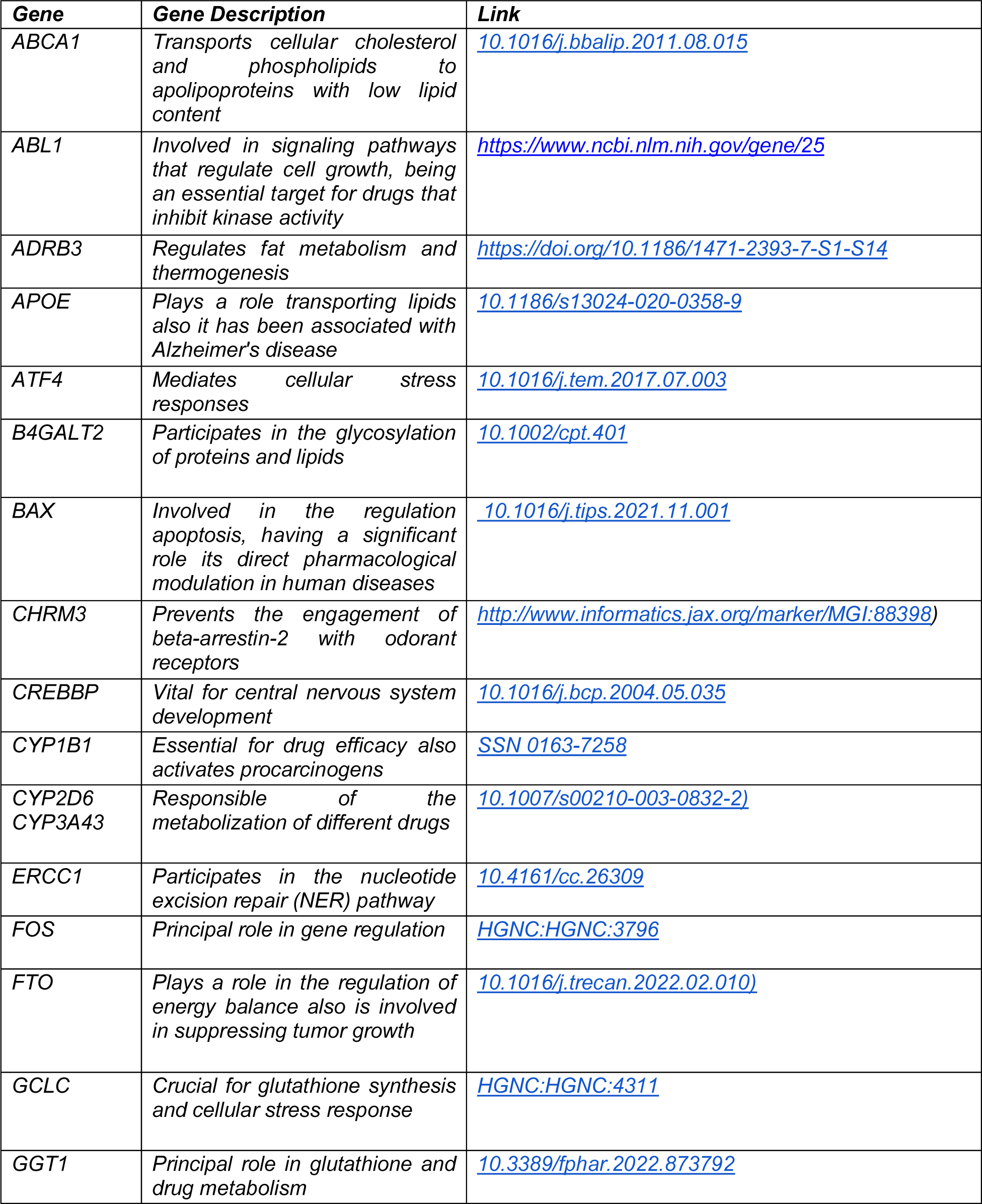
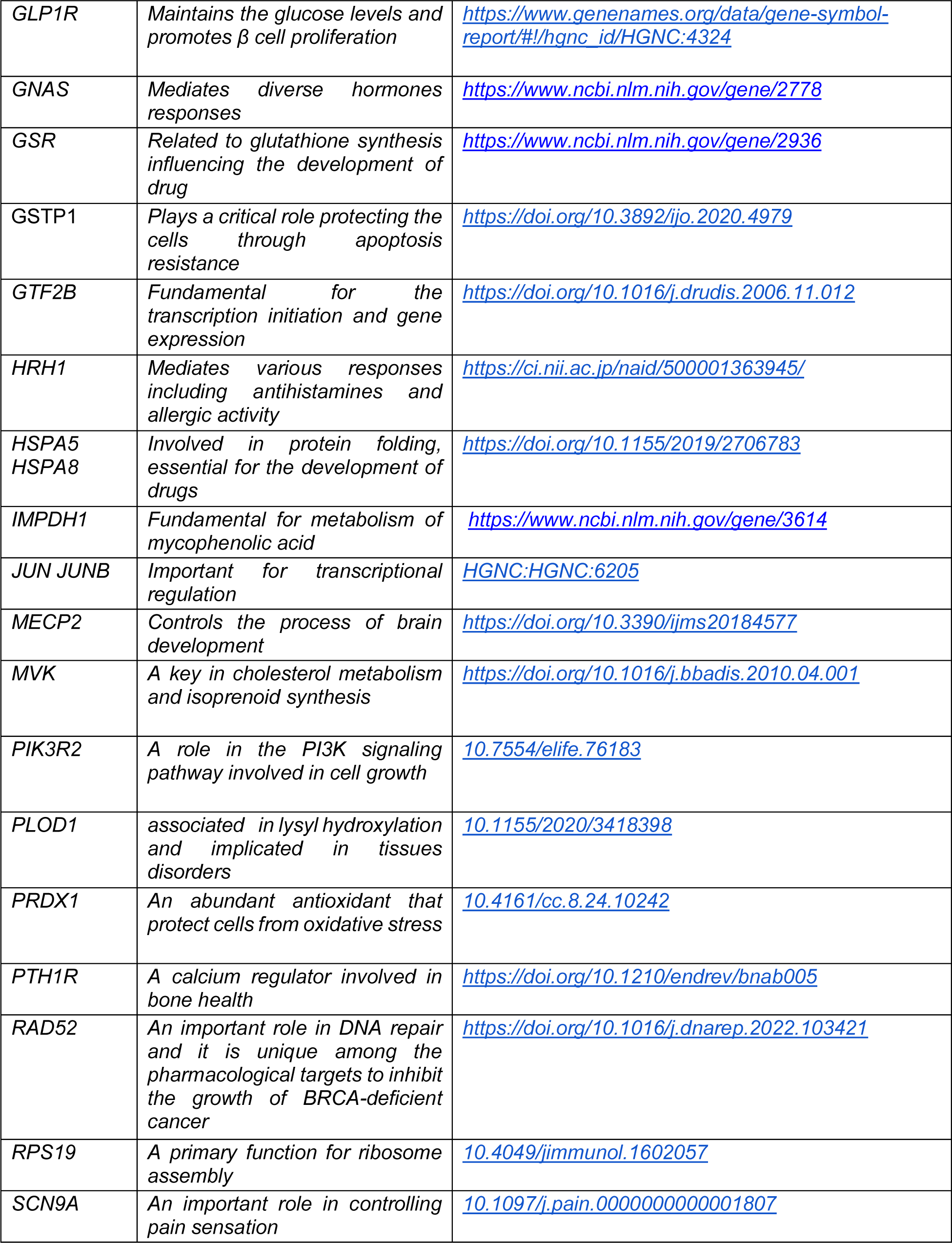
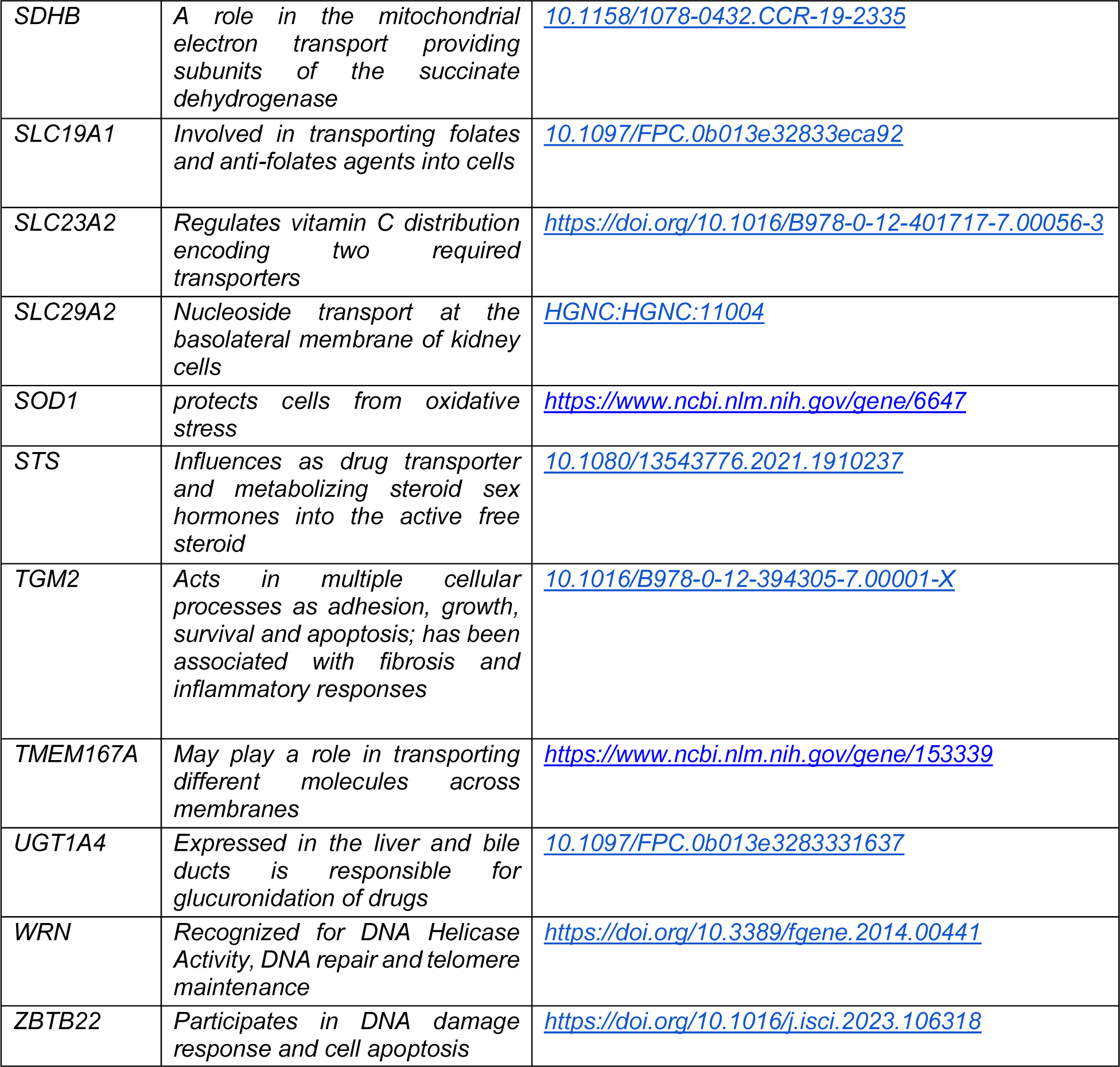
Gene Descriptions for spaceflight-flown drug associated genes.

